# Mechanism of SHP2 activation by bis-Tyr-phosphorylated Gab1

**DOI:** 10.1101/2024.12.23.630100

**Authors:** Lisa Machner, Alaa Shaikhqasem, Tobias Gruber, Farzad Hamdi, Constanze Breithaupt, Judith Kniest, Felix Wiebe, Marc Lewitzky, Christoph Parthier, Fotis L. Kyrilis, Jochen Balbach, Panagiotis L. Kastritis, Stephan M. Feller, Milton T. Stubbs

## Abstract

The non-receptor tyrosine phosphatase SHP2 (PTPN11) is a regulator of diverse cellular functions including mitogenic activation and cell migration. SHP2 consists of two tandem SH2 domains followed by the catalytic domain, and is autoinhibited by the N-terminal SH2 domain that blocks access to the active site. Mutations that influence auto-inhibition have been implicated in cancer and other diseases, and allosteric inhibitors have been developed that stabilise the inactive state. The mechanism of SHP2 activation remains unclear, however. Here, we show that the intrinsically disordered bis-phosphorylated SHP2-activating peptide pY^627^pY^659^-Gab1 binds to both SH2 domains, undergoing a partial disorder-to-order transition in the process. In addition to eliciting changes in SH2 domain dynamics, the peptide reorganises their relative orientations to provide a new SH2-SH2 interface. Our data suggest an active conformation for SHP2 that is also applicable to the hematopoietic cell-specific SHP1 (PTPN6), shedding light on the activation mechanism of both enzymes and paving the way for the development of novel compounds that modulate SHP2 activity.

## Introduction

The Src-homology 2 (SH2) domain-containing protein tyrosine phosphatase 2 (SHP2 or PTPN11) is an important regulator of cellular processes through the Ras/MAP kinase signalling pathway, which has been implicated in a number of cancers (Asmamaw et al. 2022) and is upregulated in diabetes (Saint-Laurent et al. 2022). The enzyme consists of two N-terminal tandem SH2 domains (N-SH2 and C-SH2) that bind phosphorylated tyrosine (pY) residues of binding partners, followed by a protein tyrosine phosphatase (PTP) domain [**Figure 1**]. In the absence of pY-bearing partners, the N-SH2 domain binds to the PTP catalytic site, blocking substrate access and maintaining SHP2 in an autoinhibited or “closed” state (Hof et al. 1998) [**Figure 1B**]. The crystal structure of the constitutively active oncogenic mutant SHP2^E76K^ (LaRochelle et al. 2018) reveals an open active site, with the C-SH2 domain rotated by ca. 120° with respect to the PTP domain and the N-SH2 relocated to the opposite face [**Figure 1C**]. This rearrangement can be reversed in the presence of the allosteric inhibitor SHP099 (Chen et al. 2016) to stabilize the autoinhibited state (Pádua et al. 2018; LaRochelle et al. 2018).

**Figure 1.**
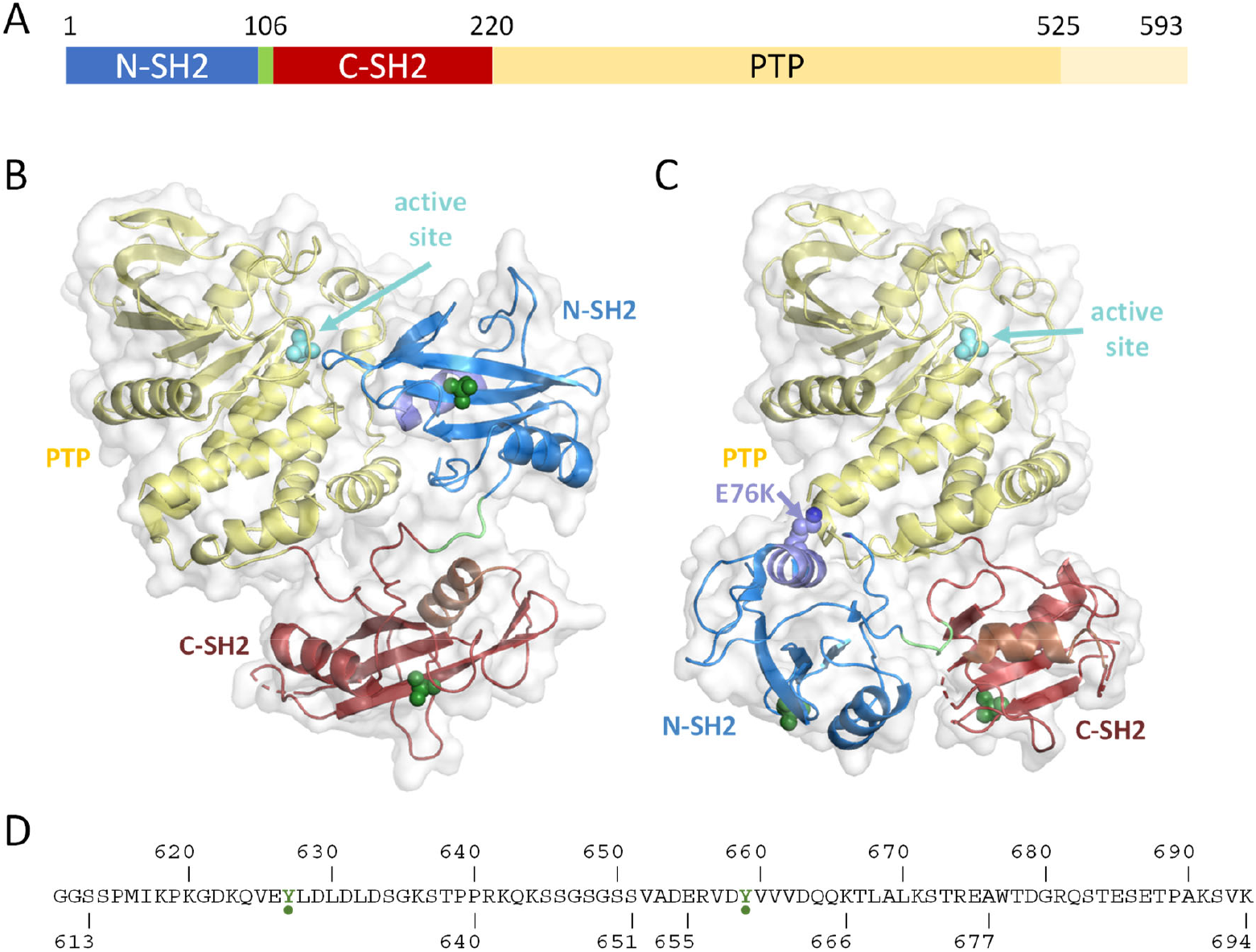
SHP2 structural elements. **(A)** Domain organisation of SHP2 (human residue numbering), consisting of two SH2 domains (blue and red) joined by a short linker (green) followed by the protein tyrosine phosphatase (PTP) catalytic domain (yellow). The latter possesses a C-terminal extension that can be dispensed with for activity measurements and is truncated in SHP2 structures. **(B)** Crystal structure of SHP2 in its autoinhibited state (Hof et al. 1998) (pdb code 2shp). The N-terminal SH2 domain blocks access to the PTP active site, marked by cyan spheres showing the phosphate position of phospho-tyrosine (pY) products. To aid orientation, locations of the SH2 pY phosphate binding sites are marked by green spheres and SH2 αB helices (see **Figure 2**) are in separate tones (αB^N-SH2^ light violet, αB^N-SH2^ brown). **(C)** The constitutively active oncogenic SHP2^E76K^ mutation E76K results in a substantial reorganisation of the N- and C-SH2 domains (LaRochelle et al. 2018) (pdb 6crf), unmasking the active site for substrate phosphorylation. Orientation of the PTP domain as in **(B). (D)** Sequence of Gab1^611-694^, with (p)Y residues (p)Y^627^ and (p)Y^659^ marked in green.

Activation must be tightly regulated, as witnessed in a variety of genetic disorders involving SHP2. 50% of patients with Noonan syndrome (NS) and 90% of patients with Noonan syndrome with multiple lentigines (NSML, formerly LEOPARD syndrome) harbour germline mutations in SHP2 (Tartaglia et al. 2001; Tartaglia et al. 2006), many of which exhibit increased basal activity (Keilhack et al. 2005). Both NS and NSML may predispose patients to juvenile myelomonocytic leukaemia (JMML), an aggressive leukaemia of infants and children, and SHP2 activation by the bacterial effector protein CagA from *Heliobacter pylori* (*Hp*CagA) has been implicated in gastric carcinogenesis (Higashi et al. 2002; Nagase et al. 2015). On the other hand, SHP2 has been demonstrated to act as a tumour suppressor in some liver and colorectal cancers (Chen et al. 2020; Bard-Chapeau et al. 2011), possibly due to SHP2-dependent dephosphorylation of the STAT family transcription factor STAT5A (Chen et al. 2003).

SHP2 is activated by pY-peptides binding to the SH2 domains, which is thought to elicit conformational changes in the enzyme (Barford and Neel 1998). Whereas some activators are high-affinity binding partners that interact solely with the N-SH2 domain (Hayashi et al. 2017; Sugimoto et al. 1994), a number of SHP2 activators possess tandem tyrosine phosphorylation sites (Ohnishi et al. 1996; Ottinger et al. 1998). Studies on the interaction of the N-SH2-C-SH2 tandem domain from SHP2 with bis-phosphorylated pYpY-peptides from *Hp*CagA (Hayashi et al. 2017) and from the programmed cell death protein 1 (PD-1) (Marasco et al. 2020) have shown that the tandem domain is able to engage both pY moieties simultaneously. The latter study revealed conformational rearrangements in the tandem domain upon pYpY-PD-1 peptide binding, which is in keeping with an opening of the active site as the short 14 amino acid pYpY-PD-1 linker (Marasco et al. 2020; Patsoukis et al. 2020) is unable to span the N-SH2 and C-SH2 pY binding sites in the autoinhibited closed SHP2 conformation (Hof et al. 1998). On the other hand, the arrangement of the SH2 domains in the constitutively E76K active open conformation (LaRochelle et al. 2018) is also incompatible with simultaneous binding of both pY moieties in PD-1, so that the mechanism whereby bis-phosphorylated peptides activate SHP2 remains unclear.

The Grb2-associated binding protein 1 (Gab1) is a hub protein composed of a folded pleckstrin-homology (PH)-domain that binds phosphatidylinositol lipids in the membrane, and a 570 aa long disordered tail that serves as a docking site for SH2 and SH3 domain-containing binding partners (Holgado-Madruga et al. 1996; Liu and Rohrschneider 2002). Phosphorylation of the two tyrosines Y^627^ and Y^659^ near the Gab1 C-terminus results in the binding and activation of SHP2 (Cunnick et al. 2001; Simister and Feller 2012). We have previously shown that the Gab1 caspase cleavage C-terminal fragment G^611^-K^694^ (Le Goff et al. 2012) [**Figure 1D**] is disordered in solution both in the unphosphorylated and bis-phosphorylated states, with residues C-terminal to (p)Y^659^ exhibiting a slight helical propensity (Gruber et al. 2022). Here, we investigate the interaction between the bis-phosphorylated fragment pY^627^pY^659^-Gab1^613-694^ and the N-SH2-C-SH2 tandem domain SHP2^1-222^.

## Results

Binding of pY^627^pY^659^-Gab1^613-694^ to the SHP2^1-222^ tandem domain was investigated initially using isothermal titration calorimetry, demonstrating a 37 ± 13 nM affinity that involves a strong enthalpic component [**Figure S1A**]. Interactions with the isolated N-SH2^1-106^ and C-SH2^102-220^ are approximately 2 orders of magnitude weaker [**Figures S1B, C**], with minor deviations from a 1:1 stoichiometry apparent from the isotherms. As the binding enthalpy to the tandem domain approximates to the sum of those for the individual domains, the interaction with the tandem domain yields a significant additional favourable entropic contribution. Mutation of the two phosphotyrosine binding sites to yield the tandem variants SHP2^1-222^-C-SH2^dead^ (R^138^A/H^169^A) and SHP2^1-222^-N-SH2^dead^ (R^32^A/H^53^A) (Hayashi et al. 2017), which in the isolated domains abrogates binding (data not shown), resulted in an expected decrease in affinity that was nevertheless higher than that for the isolated domains [**Figure S2**].

The X-ray crystal structure of the N-SH2 domain SHP2^1-106^ in complex with pY^627^-Gab1^613-651^ [**Figures 2A, S3A**] reveals binding of phosphopeptide residues pY^627^Gab1^624-632^ (amino acids -3 to +5, numbered relative to pY^627^) perpendicular to the central SH2 β-sheet in an extended (canonical) manner typical of SH2-peptide complexes (Waksman et al. 2004), with the side chains of V^625^, L^628^, L^630^ and L^632^ occupying hydrophobic surfaces of the N-SH2 domain. The binding of pY^627^pY^659^-Gab1^613-694^ to the isolated N-SH2 domain in solution was mapped using NMR [**Figure S4**]. Titration of the ^15^N,^13^C labelled phosphopeptide to the unlabelled N-SH2^1-106^ domain [**Figure S4A**] yields chemical shift perturbations (CSPs) of residues adjacent to pY^627^ in agreement with the crystal structures, as do chemical shift indices (CSIs) indicating an extended backbone conformation. Dynamic relaxation and hNOE data on the peptide (not shown) indicate conformational restriction of residues K^623^-K^636^ upon N-SH2 binding, whereas residues P^640^-V^660^ remain disordered as in the free phosphopeptide (Gruber et al. 2022). Surprisingly, CSPs are also seen around pY^659^ in the presence of the N-SH2 domain, suggesting that the latter can bind both phosphotyrosine residues. Complementary experiments titrating the unlabelled pY^627^pY^659^-Gab1^613-694^ peptide to the ^15^N-labelled N-SH2^1-106^ domain [**Figure S4B**] are also in agreement with the crystal structures. Interestingly, order parameters of ^15^N-labelled N-SH2^1-106^ in the presence and absence of pY^627^-Gab1^613-651^ reveal enhanced dynamics of the domain upon phosphopeptide binding [**Figure S4C**].

**Figure 2.**
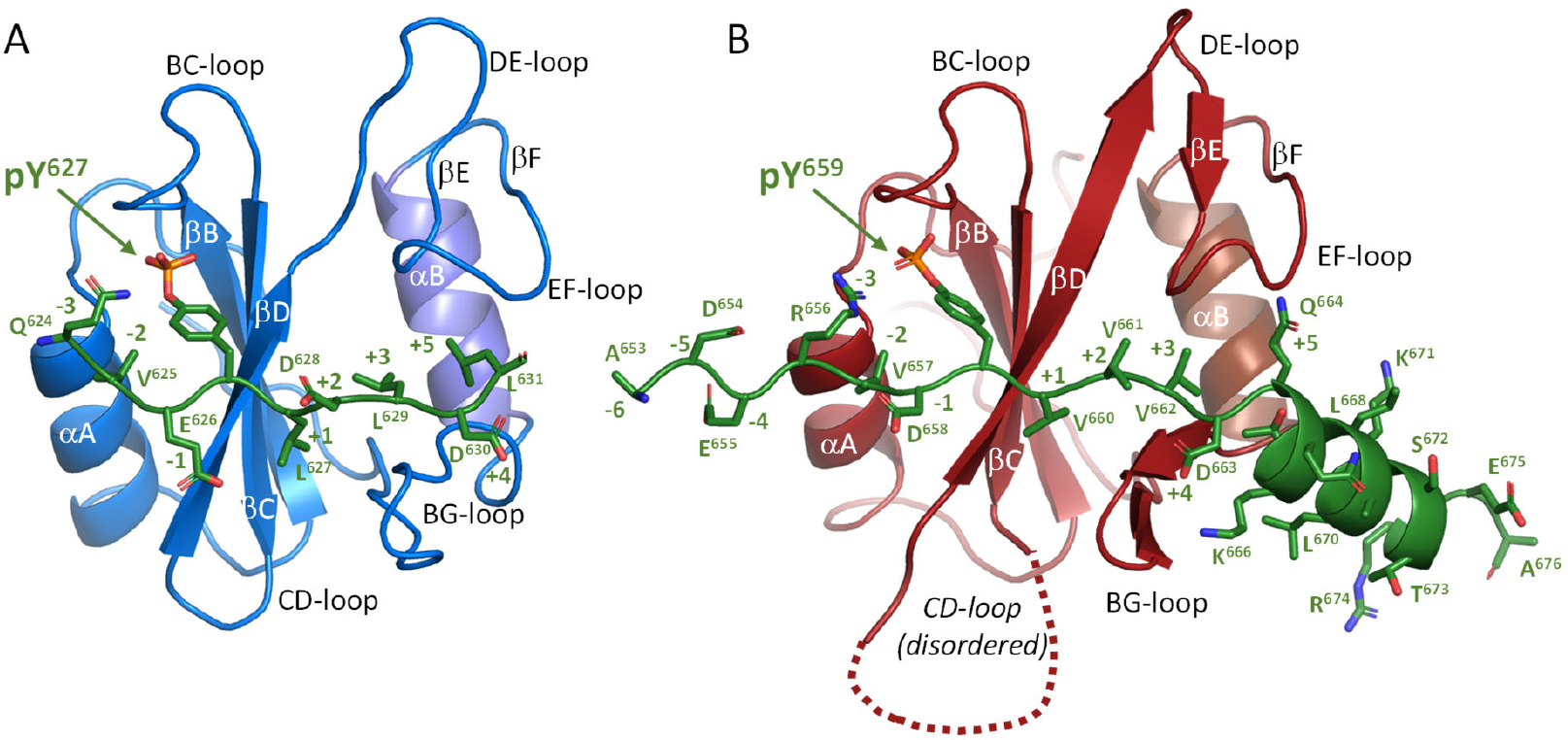
Phosphotyrosine peptide binding to SHP2 SH2 domains (see also Figure S3A). **(A)** X-ray crystal structure of the N-SH2 domain bound to the pY^627^-Gab1^613-651^ peptide (green sticks). The phosphopeptide (residue numbering from -3 to +5 relative to pY^627^) binds perpendicular to the central SH2 β-sheet in an extended manner. **(B)** Binding mode of pY^659^-Gab1^653-676^ to the C-SH2 domain determined from the electron crystallographic structure determination of the N-SH2-C-SH2-tandem SHP2^1-222^ construct in complex with pY^627^pY^659^-Gab1^617-684^; C-SH2 oriented as in **(A).** Note the hydrophilic Gln side chain in position pY^659^+5 and the presence of the C-terminal helix Q^664^-A^676^.

Although we were unable to obtain crystals of the C-SH2 domain, either in the presence or absence of peptide, the N-SH2-C-SH2-tandem SHP2^1-222^ construct in complex with pY^627^pY^659^-Gab1^617-684^ yielded long (400 µm), thin crystals with multiple lattices that proved unsuitable for X-ray crystallography. Using 3D electron diffraction (Shaikhqasem et al. 2022), however, orthorhombic microcrystals of the complex could be analysed to yield a structure to 3.2 Å resolution. Interpretable Coulomb potential density is observed for the tandem SH2 domain from R^4^ to I^221^ (with a break between T^153^ and K^164^, i.e. C-SH2 loop CD) and for the Gab1 peptide from D^622^ to S^672^, with disordered linker residues D^633^-V^652^ [**Figure S3B-D**]. Gab1 residues pY^627^Gab1^622-631^ bind to the N-SH2 domain in the canonical manner observed in the single domain structure [**Figure S3C**]. In contrast, the 24-residue segment pY^659^-Gab1^653-676^ binds the C-SH2 domain in a bipartite manner [**Figures 2B, S3D**], with amino acids -6 to +3 (A^653^ to V^662^) in an extended conformation and residues +4 to +17 (D^663^ to A^676^) forming an α-helix. The helix not only affords a larger binding interface between the C-terminal peptide and C-SH2: the canonical hydrophobic interaction between residue pY+5 (in this case Q^664^, whose side chain is disordered) with a hydrophobic patch of the signalling domain is replaced by similar interactions with the side chains of T^667^ and L^670^ (formally residues +8 and +11).

ITC experiments provide support for the unusual mode of binding to the C-SH2 domain [**Figure S5**]. Whereas titration of the monophosphorylated Gab1 peptide pY^659^-Gab1^655-677^ to C-SH2^102-220^ [**Figure S5B**] yields an affinity of 4 μM, comparable to that of the doubly phosphorylated pY^627^pY^659^-Gab1^613-694^ [**Figure S1C**], C-terminal truncation (pY^659^-Gab1^655-666^) results in a tenfold reduction [**Figure S5C**] and the corresponding C-terminal peptide Gab1^667-677^ elicits no change in enthalpy [**Figure S5D**]. NMR CSPs of the isotope labelled bis-phosphorylated peptide in the presence of the C-SH2 domain [**Figure S6A**] agree with an extensive binding site around pY^659^, comprising Gab1 residues E^655^-K^671^. The binding site is divided into two regions, with CSIs indicating residues V^661^-Q^665^ to be in an extended and residues A^669^-T^673^ in a helical conformation. Corresponding data for the labelled C-SH2^102-220^ domain with pY^659^-Gab1^655-677^ and the C-terminally truncated peptide pY^659^-Gab1^655-666^ [**Figures S6B, C**] also support the binding mode observed in the pY^627^pY^659^-Gab1^613-694^ peptide: N-SH2-C-SH2-tandem SHP2^1-222^ domain complex crystal. In contrast to the N-SH2 domain, the dynamics of the C-SH2 domain are not affected substantially by Gab1 peptide binding [**Figure S6D**].

The pYpY-Gab1-peptide-bound tandem domain exhibits a compact arrangement [**Figure 3A**], with C-SH2 P^144^ nestling in a hydrophobic pocket formed by N-SH2 residues W^6^, F^7^, F^41^, Y^63^ and L^74^ [**Figure 3B**] supported by hydrogen bonds between R^3^ and D^104^, R^7^ and E^137^ as well as N^101^ and N^219^. This is in contrast to structures of N-SH2-C-SH2-PTP SHP2 variants [**Figure 3C, D**] as well as SHP2 tandem SH2 domains in the presence of short monophosphorylated peptides [**Figure 3E-G**] that indicate the N- and C-SH2 domains are free to act independently of one another. Remarkably, a corresponding interface is found in the structure of the closely related tyrosine phosphatase SHP1 (PTPN6), an enzyme expressed primarily in haemopoietic cells, in an open conformation (Wang et al. 2011) [**Figure 3H, I**], where SHP1 C-SH2 residue P^142^ juxtaposes W^4^, F^5^, F^39^, Y^61^ and L^72^. 2D-TROSY experiments on ^15^N-labeled SHP2^1-222^ in the presence and absence of pY^627^pY^659^-Gab1^613-694^ reveal differences in the spectra in addition to those corresponding to interaction with the peptide [**Figure S7**]. Comparison of the spectra of peptide-bound ^15^N-labelled SHP2^1-222^ with those of the peptide-bound individual domains allows partial assignment of the tandem domains, and demonstrate that the interface observed in the crystal is also formed in solution upon pY-pY peptide binding [**Figure 3J**].

**Figure 3.**
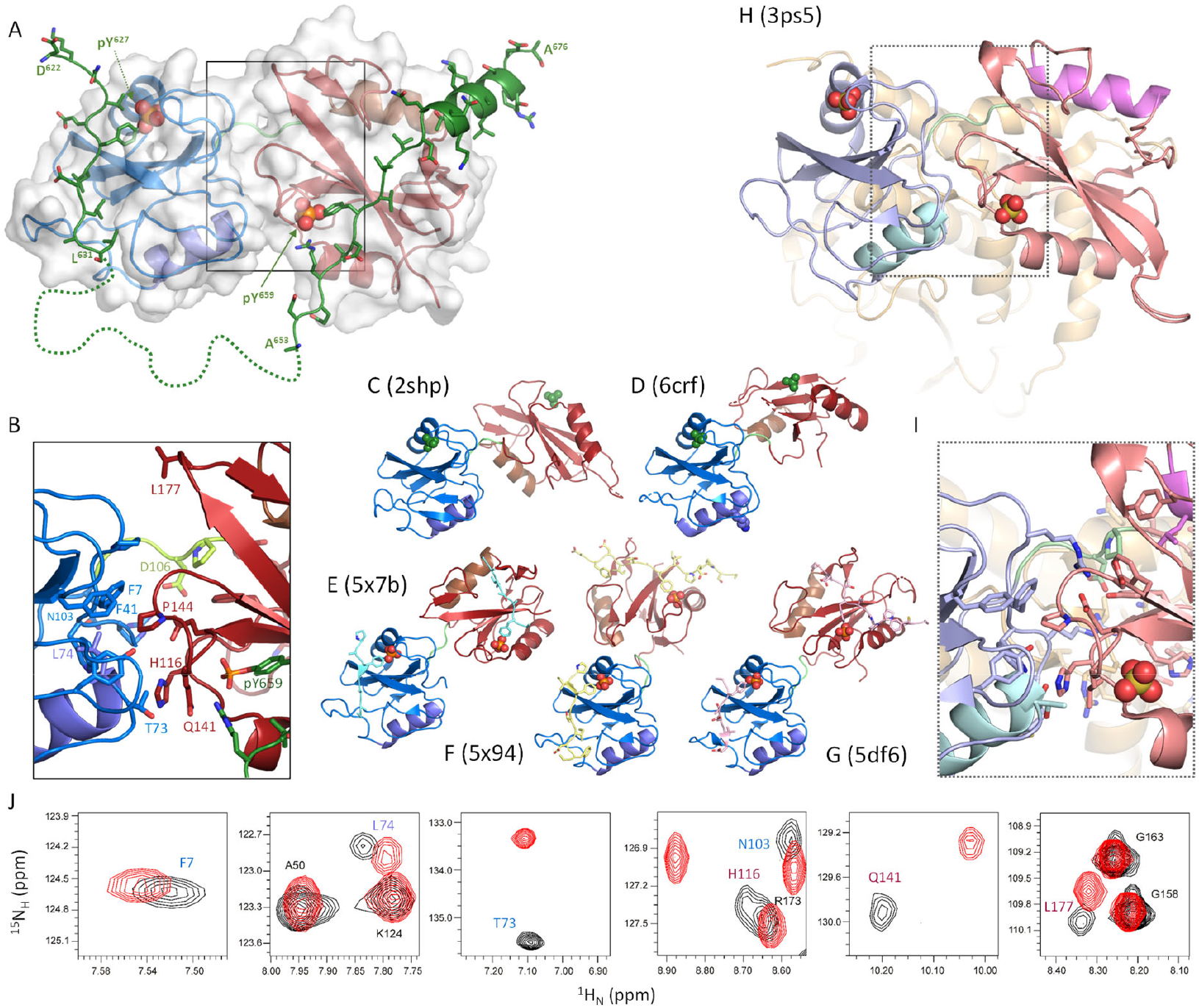
Structures of the SHP2^1-222^ tandem domain in the presence and absence of activator peptides, aligned through superposition of the N-SH2 domains (blue). **(A)** The tandem N-SH2-C-SH2 domain SHP2^1-220^ in the presence of bis-phosphorylated pY^627^pY^659^-Gab1^617-683^ (green; disordered residues Leu^632^-Val^652^ represented by a dotted line) solved by microED (this work). The crystal structure reveals a novel inter-domain interface **(B)** centred around Pro^144^ that has previously not been observed for SHP2. **(C)** SH2 domain arrangement in autoinhibited SHP2 (Hof et al. 1998), ie in the absence of any activator. Nearly all deposited SHP2 structures show this arrangement (not shown). **(D)** Relative orientations of the SH2 domains in the constitutively active oncogenic mutant SHP2^E76K^ (LaRochelle et al. 2018) without activator. **(E-G)** SHP2 tandem domains in complex with mono-phosphorylated peptides from *Helicobacter pylori* CagA (Hayashi et al. 2017) (**E, F**) or thioredoxin-interacting protein TXNIP **(G). (H, I)** An interface corresponding to that seen here **(A, B)** is found in an open crystal structure of the related SHP1/PTPN6 enzyme (Wang et al. 2011)). For orientation, SHP1 helices αB^N-SH2^ and αB^N-SH2^ are coloured cyan and pink respectively. **(J)** Selected regions of the 2D-TROSY spectra [**Figure S7**] from ^15^N-labelled SHP2^1-222^ (black) in the presence of pY^627^pY^659^-Gab1^613^-^694^ superimposed on those of the peptide bound ^15^N-N-SH2 and ^15^N-C-SH2 domains (both red).

## Discussion

The data presented here reveal details of the interaction between a bis-phosphorylated activator peptide and the N-SH2-C-SH2 tandem domains of SHP2. The pY^627^pY^659^-Gab1^613-694^ peptide binds SHP2^1-222^ with high affinity, with pY^627^ and pY^659^ binding to N-SH2 and C-SH2 respectively. Chemical shift perturbations for residues of the labelled peptide around pY^659^ in the presence of the isolated N-SH2 domain [**Figure S4A**] suggest however that both phosphotyrosine residues can bind the N-SH2 domain, providing a possible explanation for the deviation from 1:1 binding stoichiometry apparent in the ITC data [**Figure S1B**].

The N-terminal pY^627^-Gab1 peptide interacts with N-SH2 in a canonical fashion [**Figure 2A**], yet NMR data indicate that the domain exhibits enhanced dynamics upon phosphopeptide binding [**Figure S4C**]. A similar behaviour has been observed for PD-1 binding to the SHP2 N-SH2 domain (Marasco et al. 2021; Marasco et al. 2020), for which it has been postulated that pY binding perturbs a hydrogen bond network in the N-SH2 domain that propagates to the DE- and EF-loops. Ensuing conformational fluctuations in the DE-loop, which occupies the PTP active site cleft in the autoinhibited state, could result in activation of the phosphatase through disengagement of the N-SH2 domain. At the same time, opening of the EF-loop allows residues in the activating peptide C-terminal to pY to fully occupy the N-SH2 peptide binding site, preventing reclosure of the N-SH2:PTP interface. Opening of the central β-sheet of N-SH2, thought to relay the peptide-binding signal to the specificity loops, has been proposed to represent a key element of activation (Anselmi and Hub 2021).

In contrast to the canonical binding of pY^627^-Gab1 to the N-SH2 domain, the pY^659^-Gab1 peptide exhibits a bipartite binding to C-SH2. Gab1 residues adjacent to pY^659^ exhibit an extended conformation, whereas those C-terminal to pY^659^ adopt a helical conformation, where residue D^663^ could facilitate helix initiation through N-capping (Presta and Rose 1988). Such a binding mode involving concomitant peptide folding and binding may contribute to increased selectivity of the peptide for the C-SH2 domain, requiring an inherent helical propensity that is able to present the correct residues to the target domain, as has for example also been proposed for peptide recognition by class 2 G protein-coupled receptors (Parthier et al. 2007).

An unexpected finding of these studies was the presence of a novel N-SH2-C-SH2 domain interface upon pY^627^pY^659^-Gab1^613-694^ peptide binding. Additional interactions formed in the interface, accompanied by release of solvent molecules, would provide both favourable enthalpic and entropic contributions that could offset the loss of one phosphotyrosine binding site in each of the two tandem variants SHP2^1-222^-C-SH2^dead^ and SHP2^1-222^-N-SH2^dead^, explaining their higher affinity for pY-pY-Gab1 compared to either of the single domains [**Figures S1, S2**]. Notably, formation of this interface is incompatible with the closed conformation of SHP2 [**Figure S8A, B**], where the C-SH2 domain would clash with the PTP domain, yet is shared with an open state of the related tyrosine phosphatase SHP1 (PTPN6) (Wang et al. 2011) [**Figure 3A, B, H, I**], involving residues conserved between the two signalling proteins. Like SHP2, full-length SHP1 crystallises in an autoinhibited closed state [**Figure S8C**] (Yang et al. 2003) with the DE-loop of the N-SH2 domain occupying the PTP active site; the C-SH2 is in a similar location to that in SHP2 but differently oriented.

The open form of full-length SHP1 crystallised in the absence of any activators (Wang et al. 2011)), but sulphate ions from the crystallisation buffer occupy the active and SH2 phosphotyrosine binding sites [**Figure S8D**]. It has been pointed out previously that the C-SH2 domains of the oncogenic mutant SHP2 E76K (LaRochelle et al. 2018)) and the N-SH2-lacking construct SHP2 ΔN-SH2 (Pádua et al. 2018) adopt a position relative to the phosphatase domain equivalent to that in the open SHP1 structure [**Figures S8E, F**], which in turn suggests that this position of the C-SH2 domain is favoured upon disruption of the N-SH2:PTPase domain interaction. Superposition of the C-SH2 domain structure from the pYpY-Gab1-bound SHP2 tandem domain solved here upon that of the SHP2 ΔN-SH2 structure [**Figure S8G**] results in a putative full-length SHP2 domain architecture similar to that of the open form of SHP1 [**Figure S8D**]. Considering that the structures discussed here are of substantially different constructs from multiple crystal forms, we postulate that both SHP1 and SHP2 adopt a common active state [**Figure 4A; Figure S8D, G**]. Indeed, residues that make up the interface between the catalytic domain and the two SH2 domains in this conformation are conserved between the two protein tyrosine phosphatases (Wang et al. 2011)). We attribute the fact that this arrangement is not seen in the open form of SHP2^E76K^ (LaRochelle et al. 2018) to be due to the charge reversal of the E67K side chain as well as the use of a C-terminally truncated SHP2 construct for crystallisation [**Figure S9**].

**Figure 4.**
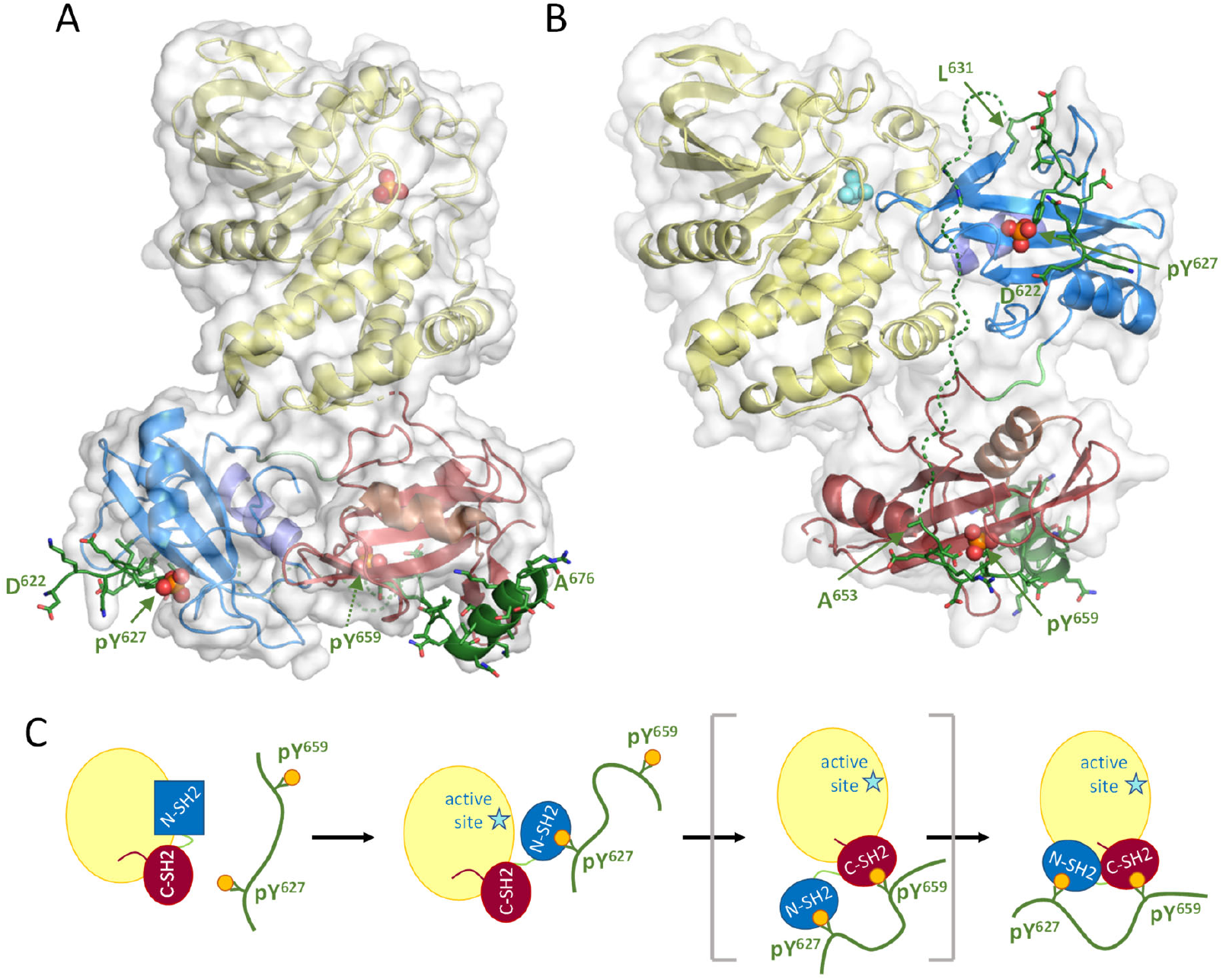
**(A)** Model of the postulated pYpY-Gab1 activated state of SHP2. **(B)** Superposition of the two phosphotyrosine peptides to the respective N- and C-SH2 domains in the autoinhibited structure requires an almost fully extended conformation to bridge by the intervening 26 residues, which would allow simultaneous binding to both SH2 domains in this conformation. **(C)** Postulated mechanism for the activation of SHP2 by bis-phosphorylated Gab1. Binding of pY^627^ destabilises the N-SH2 domain, resulting in its dissociation from the PTP domain and activation for the phosphotyrosine phosphatase. The C-SH2 domain relocates to its favoured position on the catalytic domain, engaging pY^659^, with the N-SH2 following. The multiple small energetic contributions to each state would allow for rapid conversion between inactive and active states that could be additionally modulated through e.g. dephosphorylation of the Gab1 peptide.

Availability of a structural model for the open state of SHP2 allows insights into the activation mechanism of the enzyme. Superposition of the two Gab1 phosphotyrosine peptides to the respective N- and C-SH2 domains in the autoinhibited structure [**Figure 4B**] indicates that the 26-residue pYpY-Gab1 linker (disordered in the tandem domain structure) would have to be almost fully extended in order to engage both SH2 domains in the autoinhibited state simultaneously. We envisage activation in stages [**Figure 4C**]: initial binding of the N-terminal pY^627^ peptide destabilises the N-SH2 domain, leading to its dissociation from the PTP domain.

This would allow the N-SH2 domain to dock to the C-SH2 domain to form the interface observed here, at the same time releasing the intervening Gab1 peptide residues to entropic advantage. In a final step, the SH2 domains would dock to the PTP domain to achieve the “active state” with concomitant engagement of the Gab1 C-terminal pY^659^ epitope by the C-SH2 domain. Clearly, the sequencing of these steps – some of which may occur simultaneously – is at present undetermined.

We can think of SHP2 (and SHP1) as being able to exist in two states: one closed (with a stabilised N-SH2 domain but unfavourable positioning of the C-SH2 domain) and one open (with a destabilised N-SH2 domain but stabilised N-SH2:C-SH2:PTP-domain contacts). A fine energetic balance between these states would facilitate switching between active and inactive states through ligand binding, which could be fine-tuned to achieve a wide range of signalling outcomes. In contrast to the latent activity of SHP2, SHP1 is active under basal cellular conditions, requiring down-regulation upon activation of the signalling cascade (Jones et al. 2004). This suggests that the equilibrium between active and inactive states is shifted more towards the former in SHP1, providing an explanation for the appearance of the open structure in crystals of SHP1 (Wang et al. 2011).

SHP2 activation within the cell is presumably a highly dynamic process, regulated by both Gab1 (mono and bis) phosphorylation as well as dephosphorylation. The ITC data for the N-SH2-C-SH2 tandem^dead^ domains [**Figure S2**] suggest that the transition to the compact (active) state could be induced through occupation of only one SH2 site. As such, monophosphorylated Gab1 species could elicit alternative activated species. In contrast to the 31 residue inter-pY spacing in Gab1 that requires an extended peptide to occupy both SH2 binding sites in the autoinhibited state, the insulin receptor substrate 1 (IRS-1) harbours a 49 residue inter-pY linker (Pluskey et al. 1995) that could easily allow simultaneous binding. On the other hand, the shorter inter-pY linkers found in PD-1 (14 amino acids) (Marasco et al. 2020; Patsoukis et al. 2020) and PDGFR (11 amino acids) (Ottinger et al. 1998) would preclude concurrent binding of the two phosphotyrosines in the latent state (with the caveat that the C-SH2 domain could adopt alternative positions in solution). It is therefore conceivable that the differences in linker length (as well as in amino acid composition) can result in nuanced outcomes to orchestrate cell signalling.

Much research has gone into the search for inhibitors of SHP2 due to its association with cancer, and compounds such as SHP099 (Chen et al. 2016) and TNO155 (LaMarche et al. 2020) that act allosterically by binding all three domains in the closed conformation have attracted considerable attention. The structure of the closed state has also been used to identify an alternative allosteric site at the N-SH2:PTP interface to yield SHP244 (Fodor et al. 2018), which enhances inhibition in cells when used in combination with SHP099. As SHP2 also acts as a tumour suppressor, small molecule activators may exhibit therapeutic potential (Guo and Xu 2020). Recognition of a defined active state structure paves the way towards not only a better understanding of the activation mechanisms of SHP1 and SHP2, but also the discovery of allosteric activators of these crucial enzymes that can be employed in chemical biological approaches to probe cellular functions and perhaps yield useful anticancer drugs in the future.

## Acknowledgements

This research was supported by the DFG RTG 2467 “Intrinsically Disordered Proteins – Molecular Principles, Cellular Functions, and Diseases” (project number 391498659) as well as the SFB TRR 102, DFG grants FE439/7-1 and BA 1821/6-1, the BMBF ZIK project HALOmem (grant nos. 03Z22HN23, 03COV04 and 03Z22HI2), the European Regional Development Funds (EFRE) for Saxony-Anhalt (grant no. ZS/2016/04/78115), which also provides funding for the NMR facility of the Martin-Luther-University, the Halle - Oxford (HAL-OX) research network ‘Disease Biology and Molecular Medicine’ funded by the EU/ESF and the Horizon Europe ERA Chair “hot4cryo” (project number 101086665). The authors gratefully acknowledge Dr. Ioannis Skalidis for suggesting the use of 3D electron diffraction, the success of which was central to this project, and PD Dr. Annette Meister for advice on vitrification.

## Materials and methods

### Preparation of Gab1 peptides

Cloning of the Gab1 peptide Gab1^613-694^ as a His_6_-3C-tag fusion in pET42 has been described previously (Gruber et al. 2022). Gab1^617-684^ was subcloned by cyclic polymerase extension cloning (CPEC) using the primer pair GTTCTGTTCCAGGGGCCCATCAAGCCCAAAGGAGACA / GTGGTGGTGCTCGACTCATGTGGACTGTCTCCCATC into a PCR-linearised pET42 plasmid using the primer pair TGAGTCGAGCACCACCAC / GGGCCCCTGGAACAGAAC. Both Gab1 fusion peptides were produced in *E*.*coli* and purified as described previously (Gruber et al. 2022). Briefly, fusion peptides were purified using Co^2+^ IMAC beads, processed using the 3C protease and re-applied to an IMAC column to yield the desired peptides in the flow-through. Murine ABL1 kinase was used to phosphorylate the tyrosines and the resulting phosphopeptides separated by anion exchange (HiTrap Q HP 3x 5ml). Gab1^667-677^ and the monophosphorylated peptides pY^659^-Gab1^655-677^ and pY^659^-Gab1^655-666^ were synthesised by Proteogenix, France, and the Gab1 phosphopeptide pY^627^-Gab1^613-651^ was synthesised by Davids Biotechnologie, Germany.

### Expression and purification of individual and tandem SHP2 SH2 domains

The SHP2 tandem domain N-SH2-C-SH2-SHP2^1-222^ was expressed as a His-tagged TEV-fusion protein and purified as described previously (Gruber et al. 2022). The individual SH2 domains N-SH2^1-106^ and C-SH2^102-220^ were subcloned into the pETSUMO plasmid (Invitrogen K300-01) by CPEC using primer pairs AGAGAACAGATTGGTGGTATGACCAGCCGTCGCT / CCGAATAAATACCTAAGCTTTTATTAATCTGCGCAGTTCAGCGG and AGAGAACAGATTGGTGGTCTGAACTGCGCAGATCCGA / CCGAATAAATACCTAAGCTTTTATTAGCGCGTGGTATTCAGC respectively. The vector was linearised by PCR using the primer pair TAATAAAAGCTTAGGTATTTATTCGG / ACCACCAATCTGTTCTCT. The individual domains were expressed as N-His_6_-SUMO-fusion proteins, purified using Co^2+^ affinity chromatography and cleaved overnight at 4°C using the SUMO-protease. The dialysed cleavage product was applied to a reversed-IMAC, the flow-through concentrated and further purified using size-exclusion (HiLoad 16/600 Superdex 75 pg). Corresponding procedures were used for the SH2-dead variants N-SH2^dead^, C-SH2^dead^, SHP2^1-222^-C-SH2^dead^ and SHP2^1-222^-N-SH2^dead^ (Hayashi et al 2017)), generated by QuickChange mutagenesis using the primers GATGGCAGTTTTCTGGCGGCTCCGAGCAAATCTAATCC / TCGCAATGGCGCAGTTACCGCCATTAAAATCCAGAACACG (N-SH2^dead^ and SHP2^1-222^-N-SH2^dead^) or ATGGCTCTTTTCTGGTGGCTGAAAGTCAGAGCCACC / ATGGCAAAAGCAAAGTTACCGCTGTGATGATTCGTTGTCAGG (C-SH2^dead^ and SHP2^1-222^-C-SH2^dead^).

### Isothermal titration calorimetry

ITC measurements were performed using a MicroCal^TM^ iTC200 device. All measurements were made in triplicate at 25°C in a 10 mM citrate buffer, pH 6.0 including 1 mM TCEP. All samples were dialysed in this buffer beforehand; concentrations were determined spectroscopically. Control measurements were subtracted from one-site fitted data using the Origin 7 software of the manufacturer.

### X-ray crystallographic structure determination of N-SH2^1-106^ : pY^627^-Gab1^613-651^

N-SH2^1-106^ (20 mg/ml) crystallised in the presence of a 1.5 molar excess of pY^627^-Gab1^613-651^ 15-well hanging drop plates. Tetragonal bipyramidal crystals appeared in 25 % PEG3350, 0.2 M NaCl, 0.1 M BisTris pH 5.5 within a few days.

Prior to data collection, a single crystal was flash frozen in liquid nitrogen, with 10% ethylene glycol added as cryoprotectant. Diffraction data were collected in-house at 100 K with Cu Kα radiation (λ=1.5418 Å) using a hybrid photon counting detector (HyPix-Arc 150°, Rigaku/MSC, Tokyo, Japan) mounted on a rotating anode generator (Micromax 007, Rigaku/MSC, Tokyo, Japan). X-ray diffraction data to 2.08 Å were indexed and integrated using XDS (Kabsch 2010) and the structure solved by molecular replacement PHASER (McCoy et al. 2007) using the deposited structure of N-SH2^1-106^:Gab1^621-633^ (PDB code 4qsy, Gogl and Remenyi, unpublished) as search model with the Gab1 peptide removed. The structure was completed by iterative model building in COOT (Emsley et al. 2010) and refinement using PHENIX (Liebschner et al. 2019). All structure figures were generated using PyMOL (The PyMOL Molecular Graphics System, Version 3.0 Schrödinger, LLC). Data collection and refinement statistics are listed in **Table S1**.

### Structure determination of SHP2 tandem domain SHP2^1-222^ : pY^627^pY^659^-Gab1^617-684^ *by 3DED*

Crystallisation of SHP2^1-222^ (25 mg/ml) in the presence of pY^627^pY^659^-Gab1^617-684^ (7.7 mg/ml) was performed using hanging drop vapour diffusion at 12°C in 15-well plates. Thin, needle-like crystals appeared in 29% PEG 3350, 0.1 M Tris/HCl pH 8.8 within a week. A number of crystals were washed, dissolved and subjected to SDS-PAGE, confirming the presence of two protein bands at 25 kDa and 8 kDa corresponding to Shp2^1-222^ and pYpY-Gab1^617-684^ (not shown).

Although some diffraction spots were observed to 3.5 Å on our in-house X-ray diffractometer, all macroscopic crystals exhibited severe twinning. We therefore explored 3D electron diffraction (3DED or MicroED), a continuous crystal rotation diffraction acquisition scheme (Clabbers et al. 2022). A detailed description of the procedures are provided elsewhere (Shaikhqasem et al. 2022) and are given here only briefly.

3.5 μl of the crystal sample were applied to the carbon side of a glow-discharged (PELCO easiGlow) holey carbon support film type R2/1 (200 mesh copper) Quantifoil® grid and 3.5 μl of the crystallisation buffer (diluted 1:1 with distilled water) pipetted to the back (i.e. the copper side). The grid was placed in a Vitrobot® Mark IV System (Thermo Fisher Scientific), blotted for 25 seconds at 4°C and 95% humidity with standard ashless blotting paper (Ø55/20mm, Grade 595) from the copper side, and similarly cut parafilm from the carbon side, after which it was vitrified by plunging into ethane and moved to a loading cassette.

The vitrified grid was loaded into a 200 keV Thermo Scientific Glacios Cryo-Transmission Electron Microscope and screened at low magnification microprobe TEM mode using the Falcon III EC detector to find crystals that were thinly covered in ice. Electron diffraction was detected using a scintillator-based Ceta-D camera and data collected on two crystals orientated perpendicular to each other. For each crystal, a total tilt range of 140° (-70° to 70°) was acquired, with the centre measured from -30° to 30° and wider angles measured in smaller wedges of 20° to 40° range each. The microscope was adjusted in parallel beam nano-probe mode with an exposure area limited by 50 μm C2 aperture (i.e. a beam size of 1.7 μm). For each wedge, a new spot of at least 2 µm distance was chosen on the crystal to avoid radiation damage. The electron flux was fixed to approximately 0.046 e^-^/Å^2^/s, while the total cumulative fluence was set to 3-4 e^-^/A^2^ per acquisition.

Data were processed using XDS (Kabsch 2010), yielding data to 3.2 Å with 89% completeness for the orthorhombic space group P2_1_2_1_2_1_, with one complex in the asymmetric unit. Phases were obtained via molecular replacement using PHASER (McCoy et al. 2007) as implemented in the CCP4i2 suite (Potterton et al. 2018) using the Shp2^1-222^ crystal structure in complex with a phosphorylated TXNIP-peptide (Liu et al. 2016) (PDB code 5df6) as search model, prepared as follows: phospho-peptides were deleted, and the domains were separated into the N-SH2 (residues 1-104) and the C-SH2 (residues 108-220) domains. TFZ scores were obtained of 23.1 for ensemble 1 (N-SH2) and 10.4 for ensemble 2 (C-SH2). Restrained jelly body refinements were performed with REFMAC5 (Murshudov et al. 2011) using form factors for electron diffraction. Initial model building using COOT (Emsley et al. 2010) was guided by the pre-existing structures of the N-SH2^1-106^ : pY^627^-Gab1^613-651^ complex and the C-SH2^108-220^ domain from the TXNIP complex to position the phosphorylated Gab1 peptides. After refinement, additional Coulomb potential density was visible, especially at the C-SH2 domain surface. Coordinates were further modelled into the density by hand using COOT, which was in close agreement with a Gab1 fragment generated from an AlphaFold2 (Jumper et al. 2021) model of Shp2^1-593^:Gab1^611-694^ based solely on multiple sequence alignments (Mirdita et al. 2022; Evans et al. 2021). The final model contains SHP2 residues 4 to 221 and two Gab1 fragments (622-631 and 652-672). Data collection and refinement statistics are given in **Table S2**.

### NMR spectroscopy

Labelled proteins and peptides for NMR measurements were expressed and purified as described above, with the exception of using M9 medium containing ^13^C-Glucose and/or ^15^N-NH_4_Cl.

Two-dimensional (^15^N-^1^H TROSY and ^15^N-^1^H HSQC spectra) and three-dimensional (HNCA, HNCACB, HN(CO)CACB, HNCO and HN(CA)CO) experiments were measured on a Bruker Avance III 800 MHz spectrometer equipped with a CP-TCI cryoprobe at 25°C. To measure the relaxation rates used to calculate the order parameter S^2^, additional magnetic fields were applied and an Avance III 600 MHz and a Bruker DRK spectrometer with a proton frequency of 500 MHz were also employed. Measurements were made using 500-750 µM protein sample in 20 mM BisTris/HCl, 50 mM NaCl pH 6.0 10% D_2_O (v/v) with sodium trimethylsilylpropanesulfonate (DSS) as chemical shift standard. All spectra were processed using NMRPipe (Delaglio et al. 1995) and analysed in NMRView (Johnson 2004).

^1^H-^15^N steady-state heteronuclear NOE (hNOE) experiments were measured in proton-saturated and unsaturated states. Longitudinal relaxation times (T_1_), whose inverse provides the relaxation rate 1 (R_1_ =1/T_1_), were measured by inverting the magnetisation from the z to - z-axis by a 180° pulse followed by varying delay times of 0 ms, 50 ms, 100 ms, 150 ms, 200 ms, 300 ms, 500 ms, 750 ms, 1000 ms and 1500 ms. The magnetisation was then flipped to the xy-axis by a 90° pulse for detection. To measure the transversal relaxation (T_2_), the magnetisation was excited by a 90° pulse, and a series of measurements with varying spin-lock periods of 0 ms, 8 ms, 16 ms, 32 ms, 48 ms, 64 ms, 80 ms, 96 ms, 112 ms, 160 ms were recorded. The intensity of each peak in a series of experiments was determined by NMRview or PINT (Niklasson et al. 2017).

Order parameters S^2^ were calculated from R_1_, R_2_ and hNOE measured at 800 MHz, 600 MHz and 500 MHz using the Lipari-Szabo model-free analysis (Lipari and Szabo 1982) processed in the NMR Box environment (Maciejewski et al. 2017).

Following spectra assignments of isotope labelled proteins in the free state and in the presence of a potential binding partner, chemical shift perturbations (CSPs) were calculated from the ^1^H and ^15^N chemical shift positions of corresponding residues according to the formula (Williamson 2013):

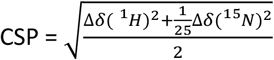

Chemical shift indices (Luca et al. 2001) for Gab1 peptides were calculated through comparison of the observed chemical shifts with a calculated chemical shift dataset for the same protein in the denatured state, taking into account phosphorylation of the tyrosine residues (Hendus-Altenburger et al. 2019):

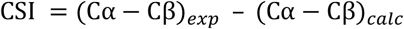

