## Supplementary Information for "Mechanism of SHP2 activation by bis-Tyr-phosphorylated Gab1"

[1]Institut für Biochemie und Biotechnologie, [2]Institut für Molekulare Medizin, [3]Institut für Physik, [4]ZIK HALOmem, Biozentrum, and [5]Charles-Tanford-Proteinzentrum, Martin-Luther-Universität Halle-Wittenberg, Halle (Saale), Germany; [6]Institute of Chemical Biology, National Hellenic Research Foundation, Athens 11635, Greece and [7]Mitteldeutsches Zentrum für Struktur und Dynamik der Proteine, Martin-Luther-Universität Halle-Wittenberg, Halle (Saale), Germany

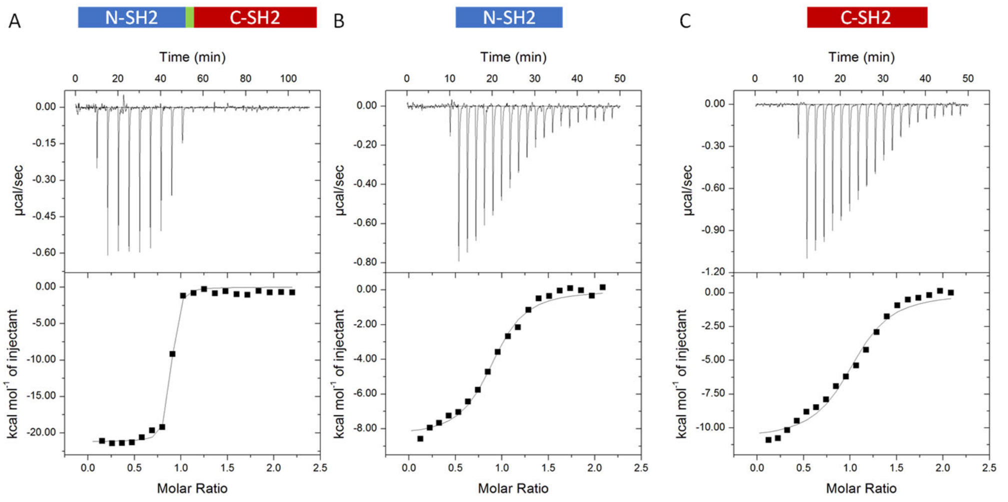


| | Shp2$^{1\text{-}222}$ : Gab1$^{613\text{-}694}$ | N-SH2$^{1\text{-}106}$ : Gab1$^{613\text{-}94}$ | C-SH2$^{102\text{-}220}$ : Gab1$^{613\text{-}94}$ |
|---|---|---|---|
| N | 0.98 ± 0.07 | 1.25 ± 0.08 | 1.14 ± 0.10 |
| $K_D$ [nM] | 37 ± 13 | 3089 ± 1881 | 2712 ± 156 |
| - ΔS · T [cal mol$^{-1}$] | 9551 ± 478 | -1508 ± 659 | 4085 ± 281 |
| ΔH [cal mol$^{-1}$] | -19743 ± 273 | -5924 ± 370 | -11677 ± 193 |
| ΔG [cal mol$^{-1}$] | -10144 ± 180 | -7432 ± 316 | -7592 ± 102 |

**Figure S1. Binding of pY$^{627}$pY$^{659}$-Gab1$^{613\text{-}694}$ to SHP2 SH2 domains using isothermal calorimetry (ITC). (A)** The tandem SH2 domain SHP2$^{1\text{-}222}$ binds with 37 nM affinity with a strong enthalpic component. **(B, C)** The same peptide binds the isolated domains N-SH2$^{1\text{-}106}$ and C-SH2$^{102\text{-}220}$ 100-fold weaker.

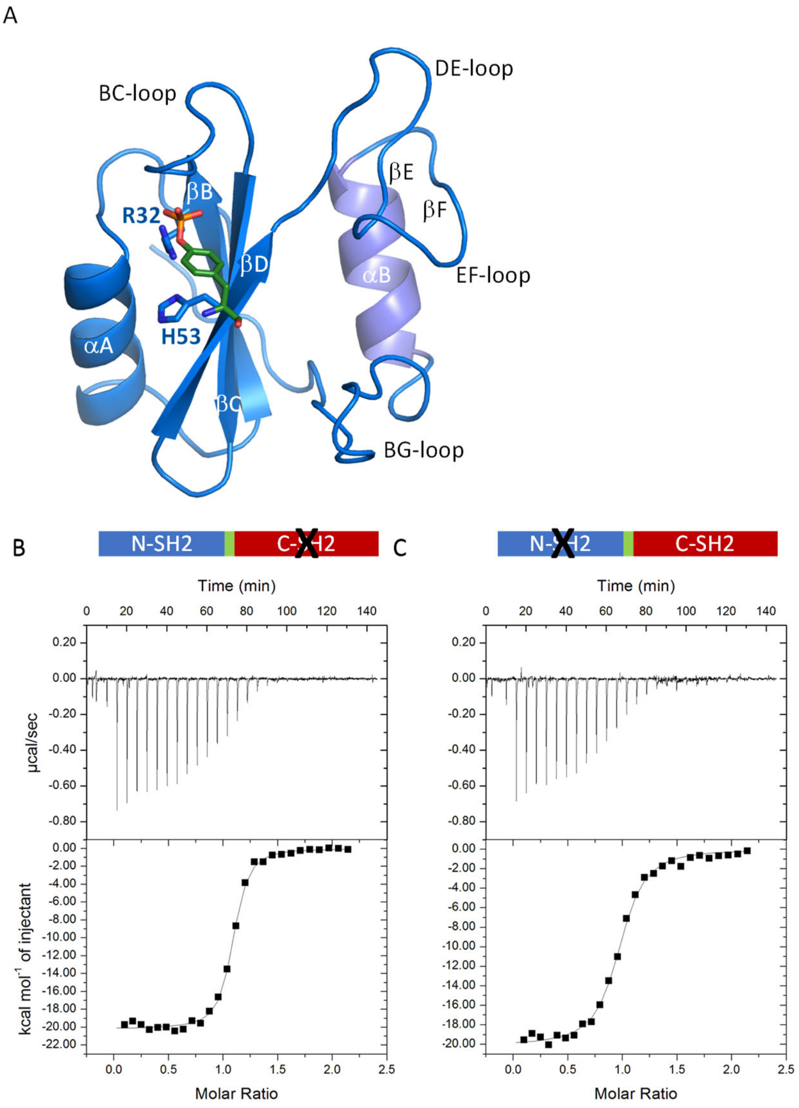


| | $Shp2^{1-222}$-C-SH2$^{dead}$ : $Gab1^{613-94}$ | $Shp2^{1-222}$-N-SH2$^{dead}$ : $Gab1^{613-94}$ |
|---|---|---|
| N | 1.04 ± 0.04 | 1.01 ± 0.07 |
| $K_D$ [nM] | 128 ± 14 | 293 ± 56 |
| - ΔS · T [cal mol$^{-1}$] | 11111 ± 394 | 11081 ± 506 |
| ΔH [cal mol$^{-1}$] | -20517 ± 345 | -19970 ± 540 |
| ΔG [cal mol$^{-1}$] | -9339 ± 52 | -8922 ± 119 |

**Figure S2. (A)** Residues $R^{32}$ and $H^{53}$ (shown in stick representation) in the N-SH2 domain, corresponding to residues $R^{138}$ and $H^{169}$ in the C-SH2 domain, constitute the binding site for phosphopeptide pY phosphate groups. Their mutation to alanine yields N-SH2$^{dead}$ ($R^{32}A/H^{53}A$) and C-SH2$^{dead}$ ($R^{138}A/H^{169}A$) (Hayashi et al. 2017). **(B)** Deleting one of the two phosphotyrosine binding sites in the tandem domain ($SHP2^{1-222}$-C-SH2$^{dead}$ ($R^{138}A/H^{169}A$) and $SHP2^{1-222}$-N-SH2$^{dead}$ ($R^{32}A/H^{53}A$)) results in reduced affinity compared to the wild type tandem domain (**Figure S1A**), yet higher than for the isolated domains (**Figure S1B, C**) with increased enthalpic and entropic contributions.

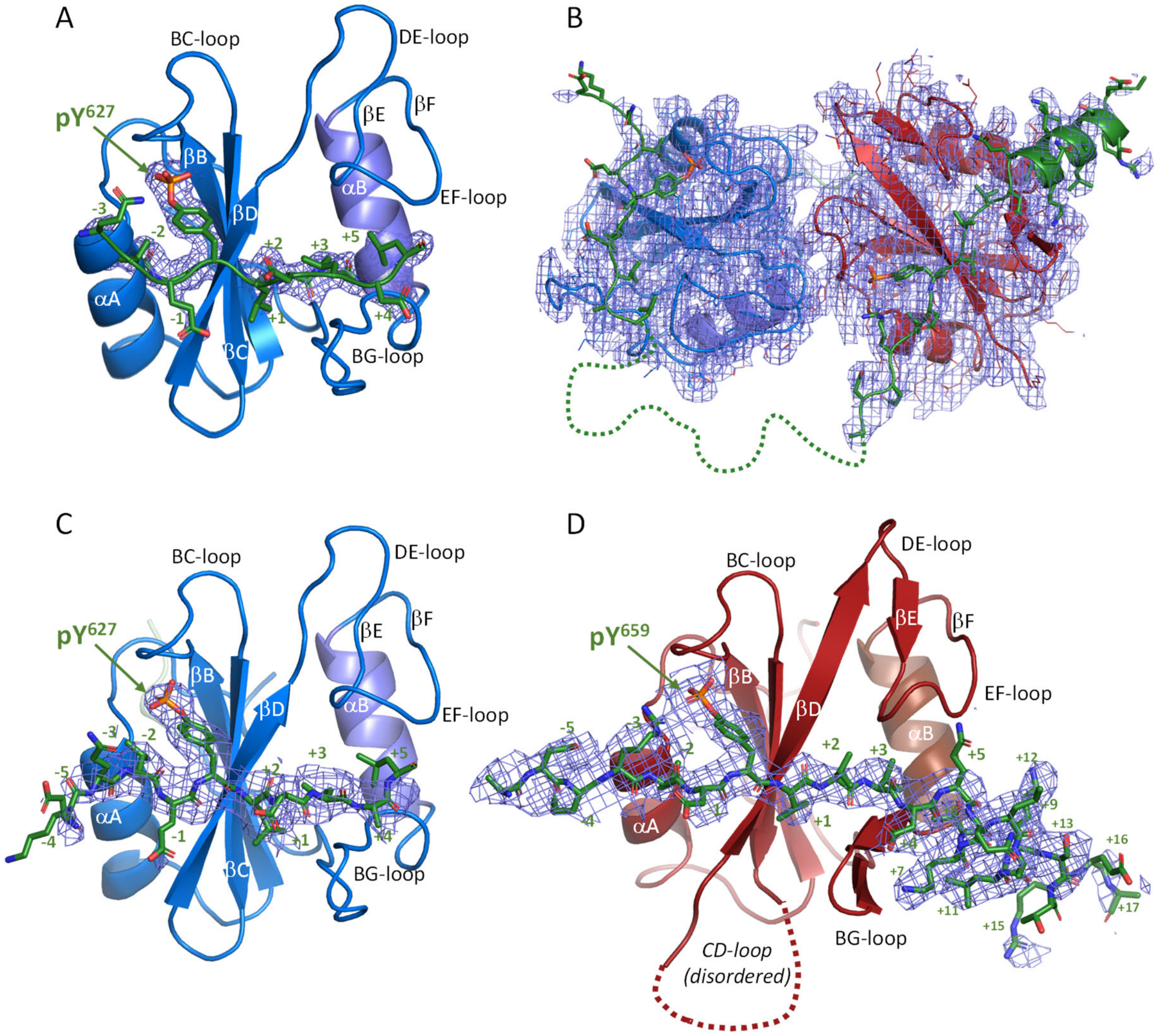


**Figure S3. Experimental crystallographic densities for phosphotyrosine peptide binding to SHP2 SH2 domains. (A)** 2Fo-Fc electron density (contoured at 1σ within 2 Å of the peptide) for the $pY^{627}$-$Gab1^{613\text{-}651}$ peptide (green sticks) bound to the isolated N-SH2 domain. **(B)** Overall Coulomb potential density (contoured at 1σ) for the N-SH2-C-SH2-tandem $SHP2^{1\text{-}222}$ construct (blue/red) in complex with $pY^{627}pY^{659}$-$Gab1^{617\text{-}684}$ (green) determined by electron crystallography. **(C, D)** Experimental 2Fo-Fc Coulomb density for the $pY^{627}pY^{659}$-$Gab1^{617\text{-}684}$ peptide bound to the SH2 domains in the tandem construct, with each domain oriented corresponding to **(A)**.

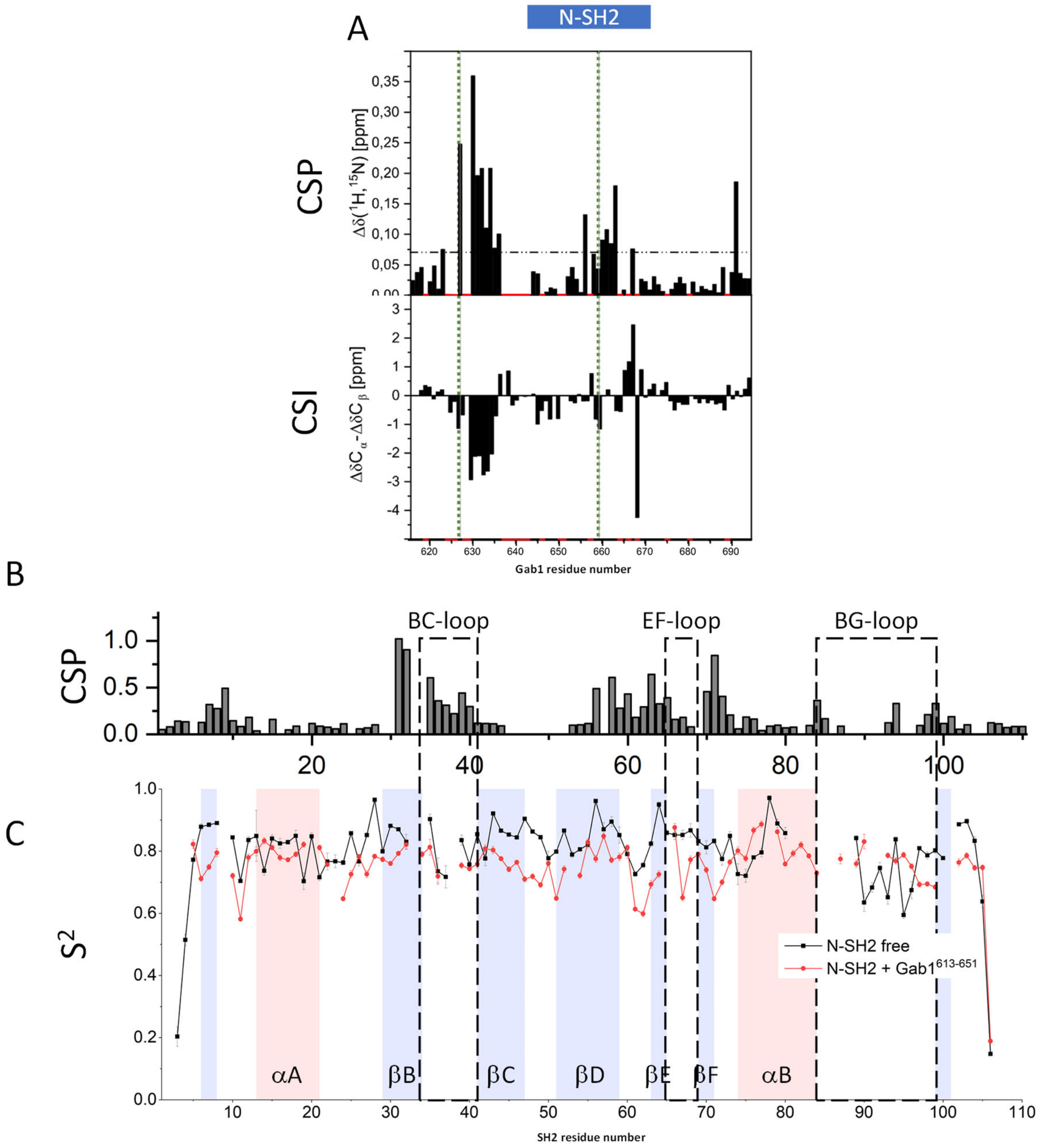


**Figure S4. Mapping the binding of pY$^{627}$pY$^{659}$-Gab1 to N-SH2$^{1\text{-}106}$ in solution. (A)** Chemical shift perturbations (CSPs) in ppm upon titration of $^{15}$N,$^{13}$C-pY$^{627}$pY$^{659}$-Gab1$^{613\text{-}694}$ to N-SH2 indicate that residues adjacent to both pY$^{627}$ and pY$^{659}$ (indicated by green dotted lines) bind to the N-SH2 domain. Chemical shift indices (CSIs), where negative values indicate extended and positive values helical backbone conformations, indicate an extended conformation for Gab1 residues 630-635. **(B)** CSPs [ppm] of $^{15}$N-N-SH2$^{1\text{-}106}$ upon binding of pY$^{627}$pY$^{659}$-Gab1$^{613\text{-}694}$ are in agreement with the X-ray crystal structure. **(C)** Per residue order parameter S$^2$ for $^{15}$N-N-SH2$^{1\text{-}106}$ for the free domain (black) and the pY-Gab1$^{613\text{-}651}$ bound complex. Secondary structure elements of the domain are indicated in blue (strand) and red (helix).

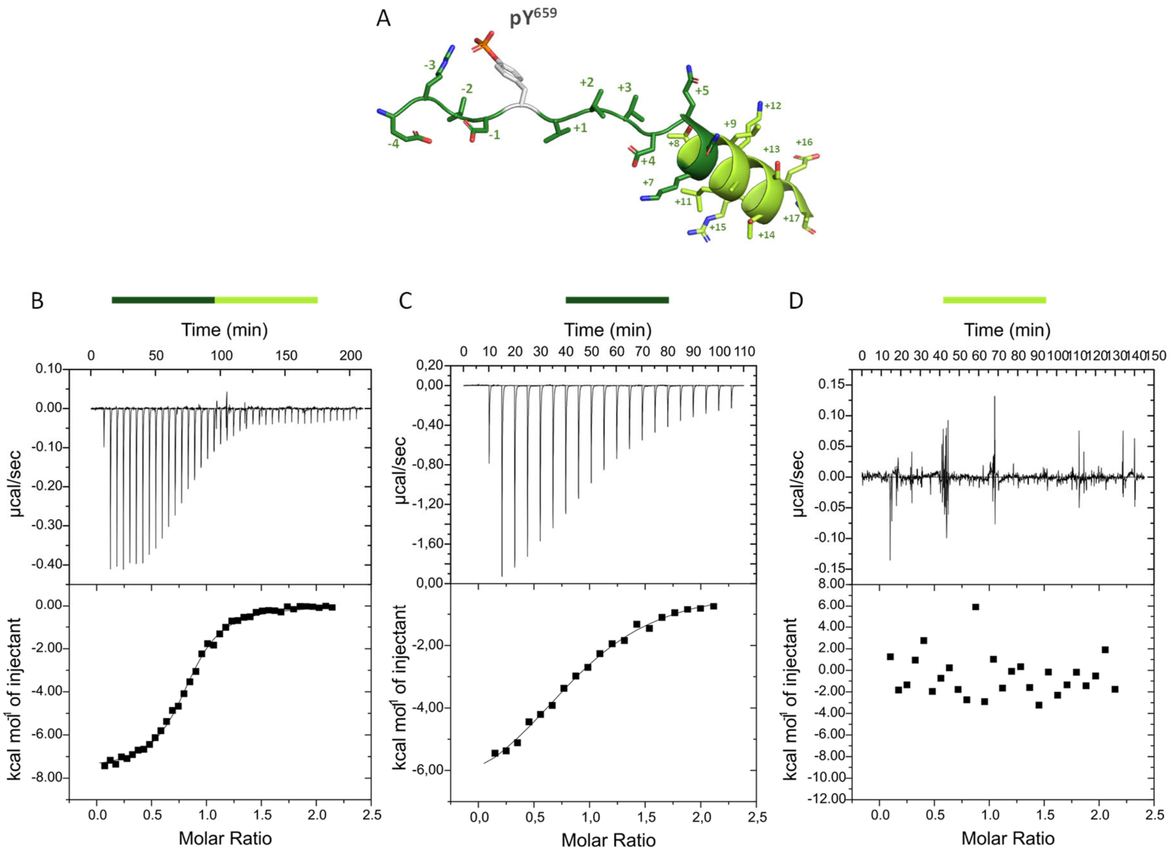


| C-SH2$^{102-220}$ binding to | | | |
|---|---|---|---|
| | Gab1$^{655-677}$ | Gab1$^{655-666}$ | Gab1$^{667-677}$ |
| N | 0.95 ± 0.11 | 1.0 ± 0.1 | - |
| $K_D$ [nM] | 4233 ± 295 | 53605 ± 8071 | - |
| - ΔS · T [cal mol$^{-1}$] | 516 ± 302 | 861 ± 129 | - |
| ΔH [cal mol$^{-1}$] | -7273 ± 631 | -6928 ± 346 | - |
| ΔG [cal mol$^{-1}$] | -7332 ± 41 | -5834 ± 85 | - |

**Figure S5. Binding of pY$^{659}$-Gab1 to the SHP2 C-SH2$^{102-220}$ domain using isothermal calorimetry (ITC). (A)** Structure of the pY$^{659}$-Gab1$^{655-676}$ peptide as bound to the C-SH2 domain in crystals of the N-SH2-C-SH2-tandem SHP2$^{1-222}$ construct in complex with pY$^{627}$pY$^{659}$-Gab1$^{617-684}$ (residues E$^{655}$-K$^{666}$ dark green, T$^{667}$-A$^{676}$ light green) **(B-D)** The monophosphorylated pY$^{659}$-Gab1$^{655-677}$ binds the C-SH2 domain with micromolar affinity, C-terminal truncation of the peptide to pY$^{659}$-Gab1$^{655-666}$ results in a tenfold drop in affinity, whereas the C-terminal peptide Gab1$^{667-677}$ shows no enthalpy change.

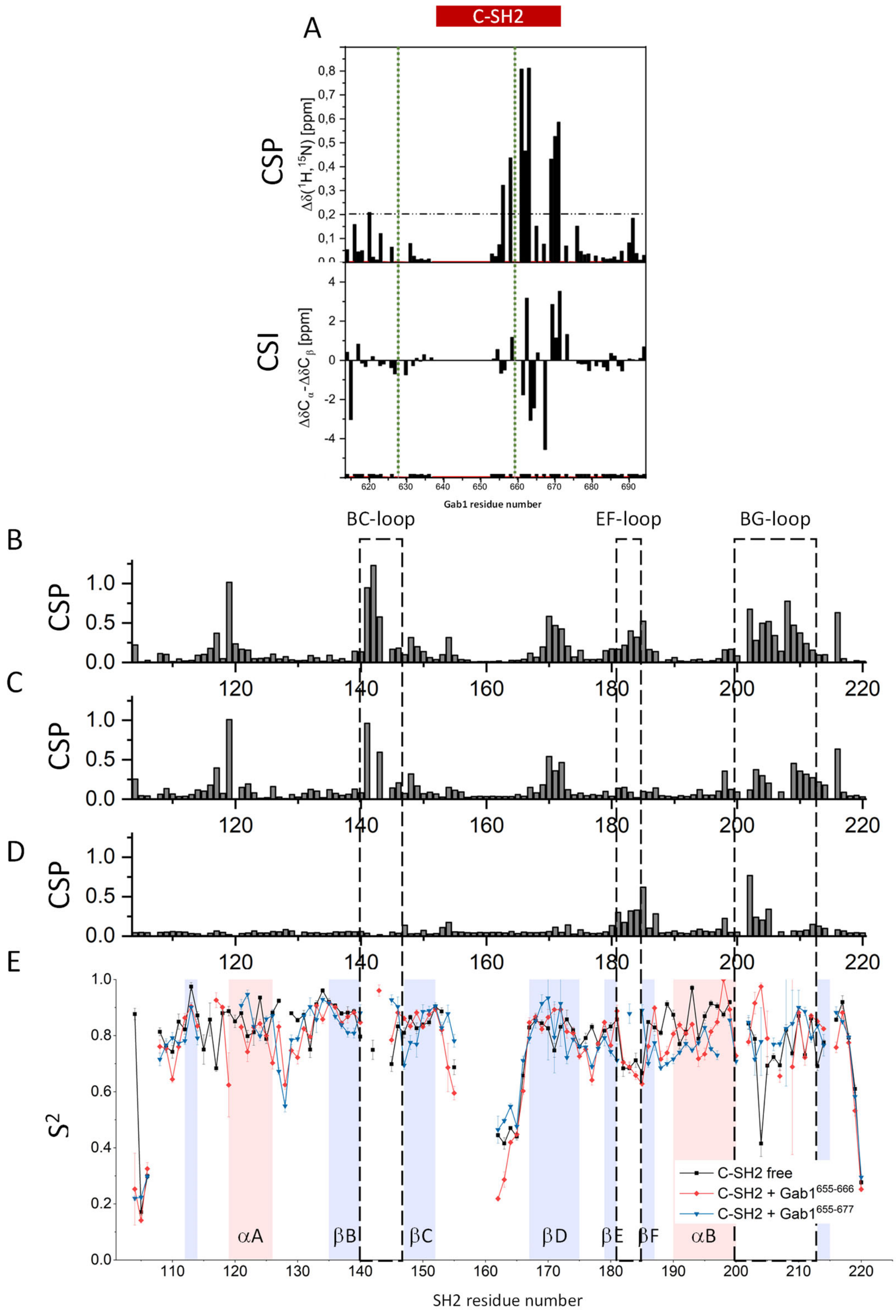


**Figure S6. Mapping the binding of pY$^{627}$pY$^{659}$-Gab1 to C-SH2$^{106\text{-}220}$ in solution. (A)** CSPs upon titration of $^{15}$N,$^{13}$C-pY$^{627}$pY$^{659}$-Gab1$^{613\text{-}694}$ to C-SH2 indicate residues 655-671 adjacent to pY$^{659}$ bind the C-SH2 domain, a sequence that is considerably longer than typical (canonical) SH2-domain binding regions. CSIs for residues 661-665 correspond to an extended conformation, whereas those for amino acids 669-673 indicate a helix. **(B)** CSPs of $^{15}$N-C-SH2$^{106\text{-}220}$ upon binding of pY$^{659}$-Gab1$^{655\text{-}677}$ are in agreement with the X-ray crystal structure. **(C)** C-terminal truncation of the peptide to pY$^{659}$-Gab1$^{655\text{-}666}$ results in reduced shifts for residues of the C-SH2 domain EF- and BG-loops. CSPs between the two bound peptides are shown in **(D)**. **(E)** Order parameter $S^2$ for the free C-SH2$^{102\text{-}220}$ domain (black), the pY-Gab1$^{655\text{-}677}$ (blue) and pY-Gab1$^{655\text{-}666}$ (red) bound complex. Residues 107, 141-144, 156-161, 201 and 215 were not assigned, and therefore no $S^2$ values could be calculated.

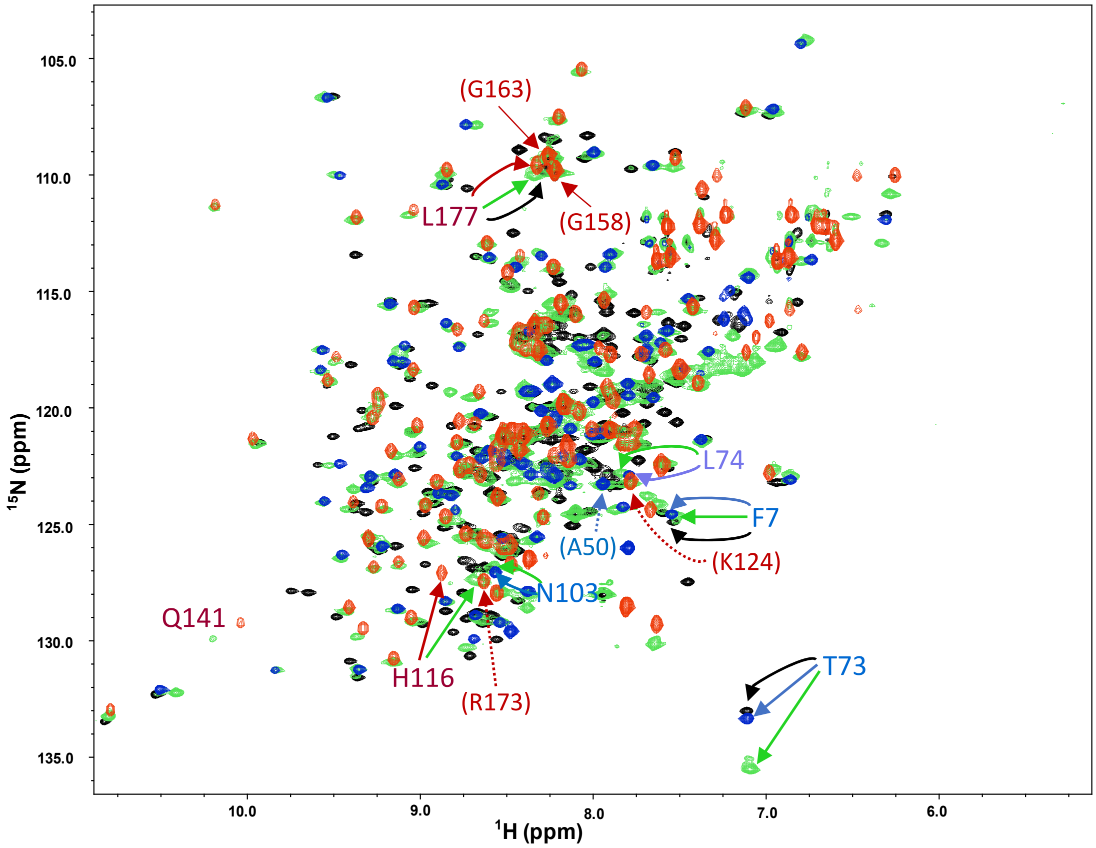


**Supplementary Figure S7.** Superposition of the $^{15}$N-TROSY spectra of the SHP2 tandem domain SHP2$^{1\text{-}222}$ in the free (black) and pY$^{627}$pY$^{659}$-Gab1$^{613\text{-}694}$ bound state (green) together with those of the bis-phosphorylated Gab1$^{613\text{-}694}$ bound $^{15}$N-N-SH2$^{1\text{-}106}$ (blue) and $^{15}$N-C-SH2$^{102\text{-}220}$ (red) domains. Residue numbers correspond to those shown in **Figure 3J**; residues in parentheses correspond to selected cross-peaks that do not shift. Note that the cross-peak for T$^{73}$ is identical in the free tandem domain (black) and the pY$^{627}$pY$^{659}$-Gab1$^{613\text{-}694}$ bound N-SH2 (blue) spectra, indicating the observed shift in the tandem domain upon peptide binding (green) is due to local change in environment in the N-SH2-C-SH2 tandem.

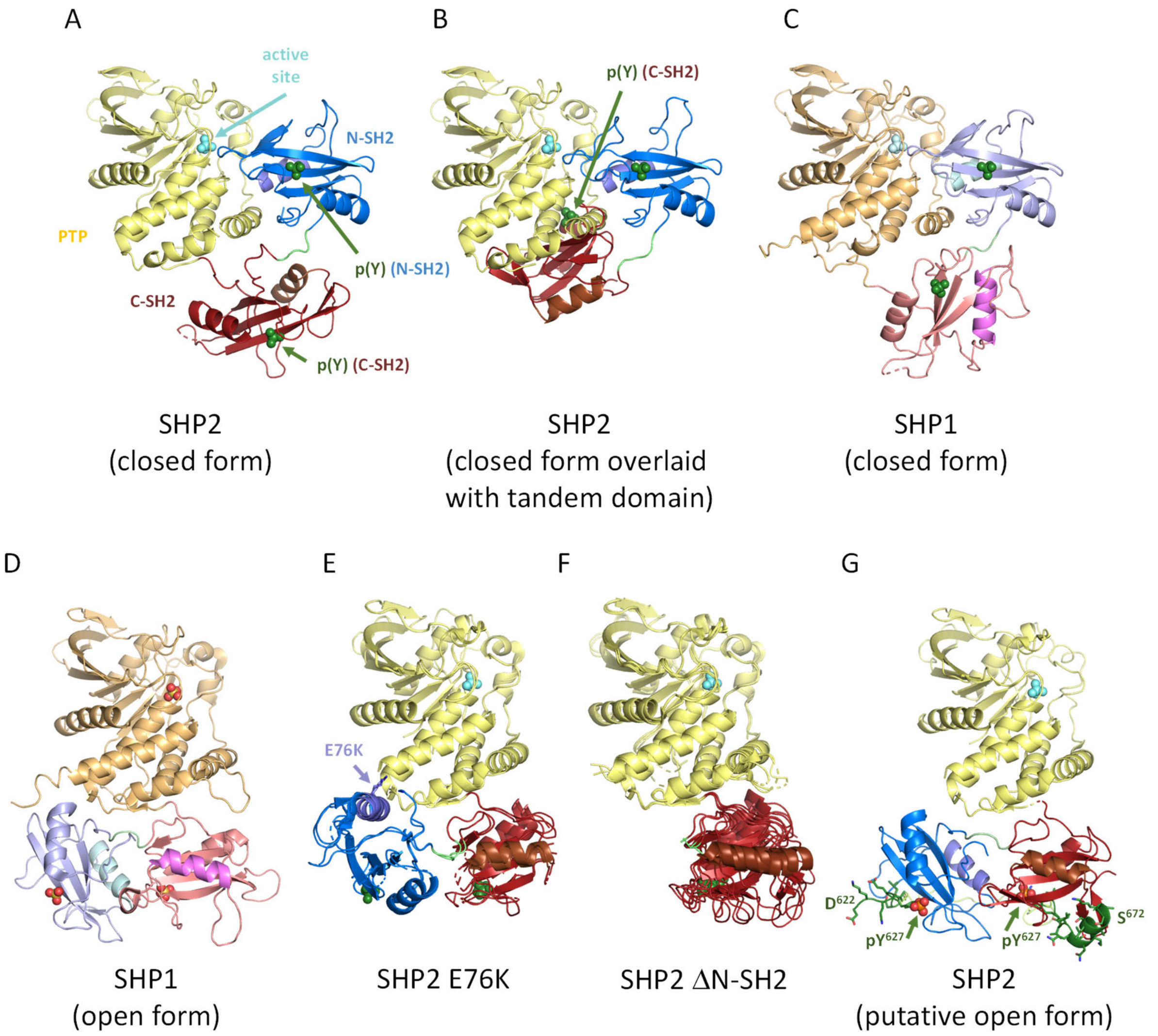


**Supplementary Figure S8. Domain architectures of SHP1 and SHP2 constructs; orientation of the PTP domain and colours as in Figure 1. (A)** Structure of autoinhibited SHP2 (pdb code 2shp). **(B)** The pYpY-Gab1-bound N-SH2:C-SH2 domain interaction seen here is incompatible with the closed conformation, as the C-SH2 domain would clash with the PTP-domain. **(C)** Structure of autoinhibited SHP1 (2b3o); colours as in **Figure 3H**. The SHP1 N-SH2 domain is in an equivalent position to that in SHP2 (**A**), but the C-SH2 is rotated. **(D)** SHP1 crystallised in an open form (3ps5) in the presence of detergent and ammonium sulphate; three sulphate ions from the crystallisation buffer occupy the PTP active site and the phosphotyrosine binding sites of the two SH2 domains (spheres). **(E)** In the absence of inhibitor, the constitutively active oncogenic variant SHP2$^{E76K}$ adopts an open conformation (6crf). In both molecules of the asymmetric unit, the C-SH2 domain exhibits the same orientation relative to the PTP domain as seen in the SHP1 open form (**D**). **(F)** Crystals of SHP2$^{\Delta N\text{-}SH2}$ lacking the N-SH2 domain (6cmq) also show the C-SH2 domain in an equivalent position in all four molecules of the symmetric unit. **(G)** Superposition of the C-SH2 domain of SHP2 tandem domain in complex with pY$^{627}$-pY$^{659}$-Gab1$^{622\text{-}672}$ on that of SHP2$^{\Delta N\text{-}SH2}$ yields an open domain architecture corresponding to that of SHP1 (**D**).

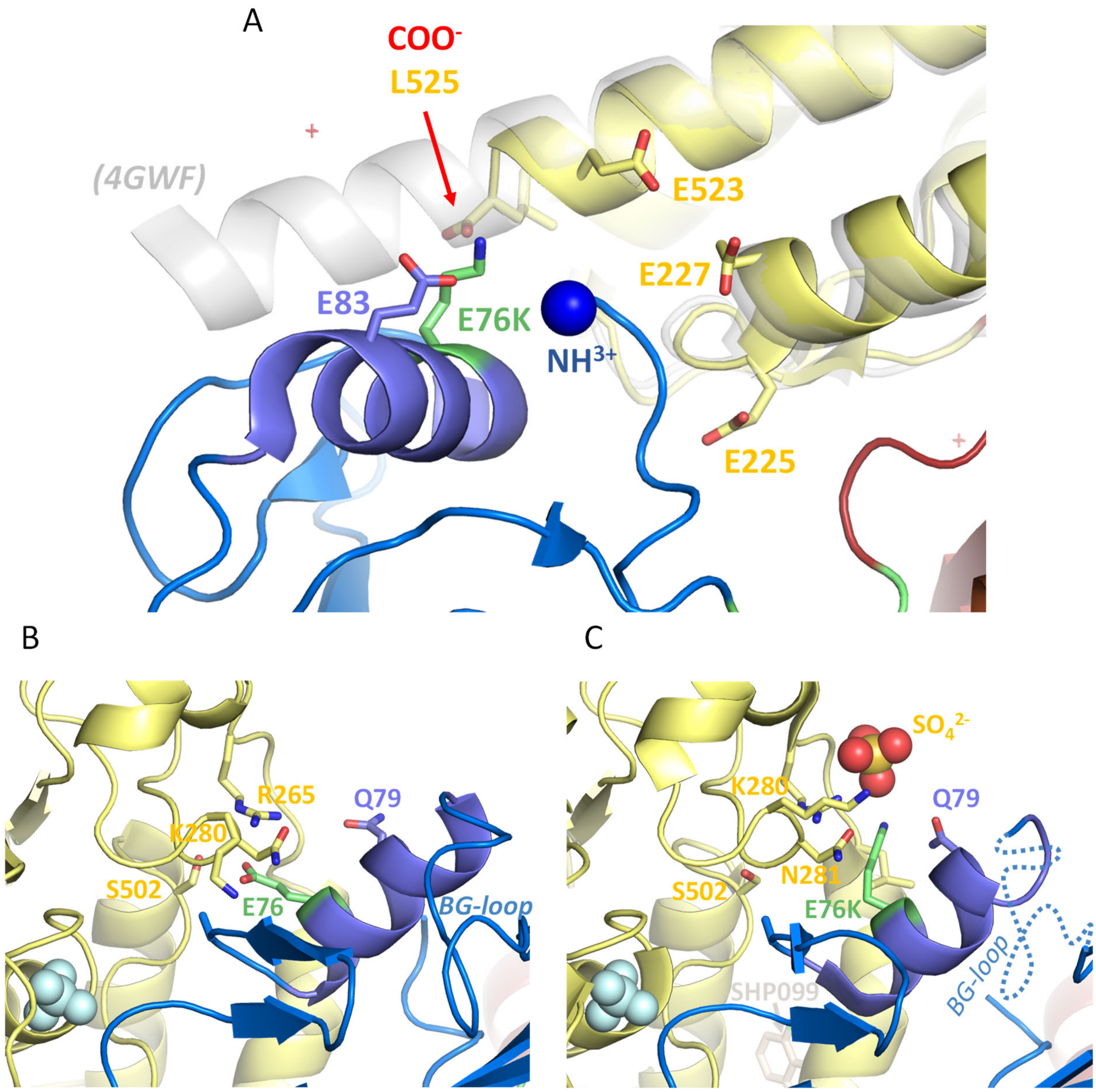


**Supplementary Figure S9. Environment of oncogenic E76(K) (A)** In the SHP2 E76K mutant structure (6crf), the position of the N-SH2 domain is perturbed due to the charge reversal of the E76K side chain, which is found together with the polypeptide amino terminus in the neighbourhood of a negatively charged patch formed by Glu83, Glu225, Glu227 and Glu523, as well as the carboxy terminus Leu525 of the SHP2 SHP2 E76K$^{1-525}$ construct used for crystallisation. The latter interaction is not available in full length SHP2; moreover, structures of SHP2 mutants using SHP2 variant$^{1-539}$ constructs (Qiu et al. 2014) demonstrate that the C-terminus extends as an $\alpha$-helix (transparent cartoon, pdb code 4gwf) that would clash sterically with the N-SH2 domain in this position. **(B)** Neighbourhood of E76 in autoinhibited SHP2 (2shp). **(C)** SHP2$^{E76K}$ adopts the closed (autoinhibited) state in the presence of the allosteric inhibitor SHP099 (6crg), but the E76K side chain is incompatible with this position, recruiting a sulphate ion from the crystallisation buffer. In this structure of the oncogenic mutant forced into the autoinhibited state, the BG loop is disordered, pointing to destabilisation of the latent state.

**Table S1: Data collection and refinement statistics for the N-SH2$^{1-106}$ : pY$^{627}$-Gab1$^{613-651}$ complex**

| | |
|---|---|
| Space group | $P4_32_12$ |
| Unit cell parameters | a = b = 60.78 Å, c = 69.85 Å<br>$\alpha = \beta = \gamma = 90°$ |
| Resolution range | 69.85 – 2.08 Å |
| Number of reflections | 118443 |
| Number of unique reflections | 8364 |
| $CC_{1/2}$ | 99.8% |
| $R_{meas}$ | 13.7% |
| $<I/\sigma(I)>$ | 14.0 |
| Completeness | 100 % |
| Multiplicity | 14.2 |
| Mosaicity | 0.29° |
| Solvent content | 38.3 % |
| Wilson B-factor | 26.4 $Å^2$ |
| Rwork / Rfree | 22.6 % / 27.3 % |
| Amino acids built:<br>SHP2 domain<br>Gab1 peptide<br>Solvent molecules | <br>99<br>9<br>43 |
| $RMS_{bond\ lengths}$ | 0.003 Å |
| $RMS_{bond\ angles}$ | 0.62° |
| Ramachandran (preferred / allowed / outlier) | 97.0 / 2.0 / 1.0 % |

**Table S2: Data collection and refinement statistics for the SHP2$^{1\text{-}220}$ tandem domain : pY$^{627}$pY$^{659}$-Gab1$^{617\text{-}684}$ complex**

| | |
|---|---|
| Excitation voltage | 200 kV |
| Wavelength | 0.025 Å |
| No of crystals used | 2 |
| Space group | $P2_12_12_1$ |
| Unit cell parameters | a = 30.59 Å, b = 82.25 Å, c = 117.76 Å<br>$\alpha = \beta = \gamma = 90°$ |
| Resolution range | 33.7 – 3.2 Å (3.39 – 3.2 Å) |
| Number of reflections | 56190 (9163) |
| Number of unique reflections | 4729 (744) |
| $CC_{1/2}$ | 94 (53) |
| $R_{meas}$ | 80.2 (220) % |
| $\langle I/\sigma(I) \rangle$ | 3.29 (1.41) |
| Completeness | 88.8 (89.7) % |
| Multiplicity | 11.9 |
| Mosaicity | |
| Solvent content | 58.5 % |
| Wilson B-factor | 47.2 Å$^2$ |
| Rwork / Rfree | 33.6 % / 39.0 % |
| Amino acids built:<br>SHP2 domains<br>Gab1 peptide<br>Solvent molecules | <br>208<br>35<br>7 |
| $RMS_{bond\ lengths}$ | 0.0019 Å |
| $RMS_{bond\ angles}$ | 1.27° |
| Ramachandran (favoured / allowed / outlier) | 95.1 / 4.5 / 0.4 % |
